# Resolving the individual contribution of key microbial populations to enhanced biological phosphorus removal with Raman-FISH

**DOI:** 10.1101/387795

**Authors:** Eustace Y. Fernando, Simon Jon Mcllroy, Marta Nierychlo, Florian-Alexander Herbst, Markus C. Schmid, Michael Wagner, Jeppe Lund Nielsen, Per Halkjær Nielsen

## Abstract

Enhanced biological phosphorus removal (EBPR) is a globally important biotechnological process and relies on the massive accumulation of phosphate within special microorganisms. *Candidatus* Accumulibacter conform to classical physiology model for polyphosphate accumulating organisms and are widely believed to be the most important player for the process in full-scale EBPR systems. However, it was impossible till now to quantify the contribution of specific microbial clades to EBPR. In this study, we have developed a new tool to directly link the identity of microbial cells to the absolute quantification of intracellular poly-P and other polymers under *in situ* conditions, and applied it to eight full-scale EBPR plants. Besides *Ca*. Accumulibacter, members of the genus *Tetrasphaera* were found to be important microbes for P accumulation, and in six plants they were the most important. As these *Tetrasphaera* cells did not exhibit the classical phenotype of poly-P accumulating microbes, our entire understanding of the microbiology of the EBPR process has to be revised. Furthermore, our new single-cell approach can now also be applied to quantify storage polymer dynamics in individual populations *in situ* in other ecosystems and might become a valuable tool for many environmental microbiologists.

## Introduction

While the demand for phosphorus (P) is strongly increasing with the growing human population, global P reserves are limited, present in only few countries, and getting increasingly more difficult to access ^1, 2^. Given the vital importance of P as a fertilizer in food production, its global scarcity is likely to become one of the greatest challenges of the 21^st^ century. On the other hand, the anthropogenic release of P is a major threat to the environment as it is a main driver of eutrophication, with major contributions from agriculture and untreated or not sufficiently treated wastewater ^3^.

Removal of P from wastewater in modern treatment plants can make an important contribution to addressing these global problems. Efficient removal of P from wastewater can prevent eutrophication in sensitive water bodies and the removed P can be applied as fertilizer ^4^. Enhanced biological phosphorus removal (EBPR) is an important biological process in wastewater treatment where P can be removed without addition of chemicals ^5, 6^. EBPR exploits the capability of certain microorganisms, termed polyphosphate accumulating organisms (PAO), to store large quantities of orthophosphate (ortho-P) intracellularly as polyphosphate (poly-P). This P-enriched biomass can be removed from the treated wastewater as surplus sludge and used directly as fertilizer or for recovery of P.

In EBPR systems, PAO are selected for by introducing alternating anaerobic-aerobic conditions ^7^. Under anaerobic conditions, PAO use poly-P as energy source to take up organic substrate and convert it to storage compounds, while under subsequent aerobic conditions, they accumulate large amounts of ortho-P from the wastewater as poly-P and respire the previously stored organic substrate. By removing biomass after the aerobic phase, poly-P can be harvested in waste water treatment plants (WWTPs). Several genera have been proposed as potential PAO, but only members of the betaproteobacterial genus *Candidatus* Accumulibacter ^8, 9^ and the actinobacterial genus *Tetrasphaera* ^10, 11^ are consistently found in high abundance in full-scale EBPR plants ^12^.

Interestingly, these two PAO occupy different ecological niches within EBPR plants. *Ca*. Accumulibacter, for which no pure culture is available, have been intensively investigated in lab-scale enrichment cultures with *in situ* and meta-omic based expression studies to verify their proposed physiology ^13, 14, 15, 16^. During anaerobic periods, *Ca*. Accumulibacter takes up volatile fatty acids (VFAs), such as acetate, and stores them as intracellular polyhydroxyalkanoates (PHAs), with the expenditure of energy generated from the hydrolysis and release of intracellular poly-P reserves. Hydrolysis of glycogen stores provides additional energy and the reducing power required for PHA storage ^17^. In subsequent aerobic periods, intracellular PHAs are respired to provide energy for cellular metabolism, glycogen generation as well as ortho-P uptake to replenish the poly-P reserves ^17^. Species in the genus *Tetrasphaera* are much less studied and their physiology is still poorly understood. Members of the genus have a diverse physiology that includes aerobic respiration, denitrification and fermentation. In the absence of oxygen, aerobically stored polyphosphate provides an additional energy source ^17, 7^. Annotation of representative genomes for the genus suggests that glucose could be stored as glycogen (but not PHAs) under anaerobic conditions, and some un-polymerised fermentation products and amino acids have been shown to accumulate in the cell. Under subsequent aerobic conditions, glycogen and accumulated intracellular substrates are suggested to be utilized for growth and the replenishment of poly-P reserves ^18; 19^.

Despite the global relevance of the EBPR process, the actual contributions of *Ca*. Accumulibacter and *Tetrasphaera* sp., respectively, to bulk P removal are still unknown as no technique was available for quantifying clade-specific contributions to the EBPR process. For such measurements not only the *in situ* abundances of PAO clades must be determined, but also quantitative *in situ* determinations of their intracellular P content at the different stages and conditions of the EBPR process are necessary. Poly-P is difficult to quantify and is usually analyzed after extraction ^20^, which makes it impossible to quantify the poly-P content in specific microbial populations. Instead microscopy-based single cell methods are needed, such as staining of poly-P using 4',6-diamidino-2-phenylindole (DAPI) ^21^ or Neisser ^8^. However, these staining methods have only been applied to yield qualitative information or relative quantitative estimates and their specificity towards poly-P is questionable - e.g. the DAPI – RNA complex is known to interfere with poly-P fluorometric quantification ^20^.

During the last decade, Raman microspectroscopy has increasingly been used to investigate physiological features of individual microbial cells in complex environmental samples ^22, 23, 24, 25^. In addition to single cell isotope labelling studies, the identification and quantification of intracellular storage polymers by this vibrational spectroscopic technique can provide useful information for microbiologists ^26, 27, 28^. Raman microspectroscopy has recently been applied to monitor glycogen, polyhydroxyalkanoates (PHA), and poly-P in randomly selected microbial cells from EBPR systems ^29, 30, 31^, but has not yet been used for absolute quantification of these storage polymers. Furthermore, simultaneous identification of the Raman-analyzed cells in EBPR plant biomass by fluorescence *in situ* hybridization (FISH), which is necessary to assign *in situ* storage patterns to specific PAO clades, has not yet been performed.

In this study we developed a Raman microspectroscopy-based quantitative approach to determine the levels and dynamics of intracellular polyphosphate and other storage polymers (PHA and glycogen) in FISH-identified *Tetrasphaera* and *Ca*. Accumulibacter cells. This approach was applied to reveal which of these PAO is most important for P removal in eight full-scale EBPR treatment plants and to validate the suggested models of their ecophysiology in these systems.

## Materials and Methods

### Bacterial strains and activated sludge sampling

The activated sludge isolate *Tetrasphaera elongata* strain Lp2 (DSM 14184) was grown aerobically in modified R2A medium (starch and sodium pyruvate were excluded and glucose was main substrate) at 26°C. The *Acinetobacter junii* culture (DSM 14698) that was used for investigating cell fixation effects on intracellular poly-P content was grown on nutrient agar (beef extract 3 g l^-1^ peptone, 5 g l^-1^, agar 15 g l^-1^).

Eight different WWTP were investigated in this study. Details about their design, operation, and performance are given in Table S1 and S2. All plants had stable P removal in the sampling period with average effluent P concentrations below 1 mg P l^-1^. All plants had minor addition of iron salts to support the EBPR process (molar ration of Fe-dosage/incoming total P of <0.5). For P uptake/release experiments, activated sludge samples from Aalborg West WWTP were collected from the aeration tank and transported at 4°C to the laboratory. For quantification of the relative abundance and the cell-specific content of P inside PAO directly in full-scale EBPR plants, sludge samples from the anaerobic (An), hydrolysis (Hyd), anoxic and denitrifying (DN), oxic and nitrifying tanks (N), and return activated sludge (RAS) were immediately fixed *in situ* in either 96% ethanol: 1 x phosphate buffered saline – 1:1 (for Gram-positive cells) or 4% PFA (for Gram-negative cells) fixative solutions ^32^. They were homogenized in a mechanical homogenizer (300 RPM, 10 min, Buch & Holm – Heidolph, Germany) in order to disrupt large sludge flocs. Samples were further homogenized in ultrasonic homogenizer bath (Branson Ultrasonic 5800, USA) at 40 kHz for 5 min. The settings of the homogenization procedures used were too weak to interfere with cell integrity ^33^. Homogenized samples were aliquoted onto CaF_2_ Raman windows and air dried for later FISH-Raman analyses.

### Phosphorus uptake-release experiments in *T. elongata*

*T. elongata* cells were harvested by centrifugation (4,500 x g, 15 min) and washed twice with chemically defined, modified mineral salts-vitamin medium (MSV) as described by ^34^. Harvested cells were re-suspended in MSV and aerobically incubated for 4 h to exhaust all intracellular carbon sources. The cultures were then made anaerobic by purging nitrogen gas for 15 min in sealed serum vials and supplemented with 200 mg l^-1^ COD equivalents of glucose as the sole carbon source. The cultures were kept anaerobic for 3 h with shaking and the cells were harvested and washed with MSV to remove residual substrate and secreted fermentation products. The cultures were then incubated under aerobic conditions for 3 h with shaking and addition of 0.5 mM phosphate. A final concentration of 0.32 mM Na_2_HPO_4_ and 0.18 mM NaH_2_PO_4_ was added as a P source and as a buffer (to retain pH at around 7.2), but no carbon sources were provided. The culture was sampled at set time intervals and the experiment was conducted in duplicate. The control experiments included heat-killed *T. elongata* biomass (autoclaved at 121°C/15 min) and a living culture with no glucose supplemented during anaerobic incubation.

### Activated sludge batch experiments for phosphorus uptake and release

Batch experiments were conducted on fresh activated sludge to analyse the P-content per cell of FISH-identified *Tetrasphaera* or *Ca*. Accumulibacter under anaerobic conditions (low P content expected) and under aerobic conditions (high P content expected). For this purpose, fresh sludge was diluted to a total suspended solids concentration of 1 g l^-1^ using filtered (0.22 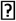m, Whatman, UK) effluent and aerated for 1 h to exhaust most intracellular carbon source reserves. Acetate and glucose were then added to a concentration of 1 mM and 2 mM, respectively, and the sludge was further supplemented with casamino acids to a COD equivalent of 50 mg l^-1^ (final COD ca. 500 mg l^-1^). The amended sludge was then made anaerobic by purging with nitrogen gas for 15 min in serum vials and was kept at room temperature (~22°C) with shaking for 4.5 h. Subsequently, the biomass was spun down (4,500 x g, 15 min) and the supernatant was discarded to remove unused substrates. The biomass was washed twice with filtered (0.22 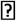m, Whatman, UK) wastewater. In the next step, 0.5 mM phosphate (15.5 mg P l^-1^) was added and the sample was made aerobic by purging with compressed air. The sludge was kept aerobic with shaking for 4.5 h without supplementation of a carbon source. pH was 7.0–7.2 throughout the experiments. Samples were taken every 30 min during both anaerobic and aerobic stages and the experiment was conducted in duplicate. Control experiments were performed with living biomass without supplementing organic substrate and with heat-killed (autoclaved, 121°C/15 min) biomass. Activated sludge samples were fixed, homogenized and aliquots air dried on CaF_2_ Raman windows for later FISH and Raman analyses.

### Raman micro-spectroscopy and calibration of the instrument

A Horiba LabRam HR 800 Evolution (Jobin Yvon, France) Raman micro-spectrometer instrument equipped with a Torus MPC 3000 (UK) 532.17 nm 341 mW solid-state semiconductor laser was used for all experiments. The laser power was attenuated to 2.1 mW μm^-2^ incident power density on the sample by a set of neutral density (ND) filters. An integrated Olympus (model BX-41) fluorescence microscope was used for selecting FISH probe labelled cells of interest for Raman analysis. A dry objective with a numerical aperture of 0.75 (Olympus M Plan Achromat, Japan) (corresponding to a measured laser spot size of about 2.6 μm – Supplementary text-1) and a magnification of 50X (working distance 0.38 mm) was used to focus the laser beam on the sample and to collect the Raman scattered light. According to the Rayleigh criterion, these optical attributes of the system translate to a spatial resolution of approximately 0.5 μm and an axial resolution of about 1.4 μm. The grating used during all measurements was 600 mm/groove and the spectral range chosen spanned from 200 cm^-1^ to 1850 cm^-1^. The wavenumber region from 300 cm^-1^ to 1800 cm^-1^ is known as the fingerprint region in terms of characterisation of biological material, as it contains the most important spectral features^35^. The Raman spectrometer slit width was 100 μm and the Raman confocal pinhole diameter was 72 μm during all measurements. Raman scattered light was detected by an Andor Charge Coupled device (CCD) (UK) cooled at −68°C. The spectra were recorded and processed using the LabSpec version 6.4 software (Horiba Scientific, France). All spectra were baseline corrected using a 6^th^ order polynomial fit and the cosmic ray interferences were removed using the cosmic ray removal feature in the software. All spectra were averages of two individual spectra with 10 s integration time. The Raman spectrometer was calibrated before all measurements to the first-order signal of a silicon wafer occurring at 520.7 cm^-1^. All Raman sample measurements were conducted on optically polished CaF_2_ windows (Crystran, UK), which produces a single strong Raman signal at 321 cm^-1^, that also serves as an internal reference point in every spectrum.

### Raman spectroscopy-based quantification of polyphosphate

The method used for quantification of polyphosphate in this study relies on a linear dependence of the Raman signal on the amount of analyte (poly-P) per unit surface area in the sample. For all cellular measurements, it was assumed that the cell monolayers were transparent and had a negligible absorption. Poly-P is a strong Raman scatterer with characteristic signature peaks occurring at 690 cm^-1^ and 1170 cm^-1^ wavenumber regions, attributed to –P-O-P- stretching vibrations (phosphoanhydride bonds) and Po_2^-^_ stretching vibrations ^29^ (Fig. S1), respectively. Both peaks could be observed simultaneously and unambiguously at the correct wavenumber region, in all of the poly-P containing spectra. Furthermore, the Raman signature band at 1170 cm^-1^ of poly-P is not found at relevant intensities in spectra from other phosphate containing cellular chemical species such as ATP and AMP (Fig. S2). The strongest Raman peak for poly-P appears at 1170 cm^-1^ in the spectrum, whereas the strongest Raman peaks for ATP and AMP occur at 1125 cm^-1^ and 990 cm^-1^, respectively.

The material density of poly-P standard solutions mounted and dried on CaF_2_ windows exhibited a linear correlation to the Raman signal intensity at the wavenumber 1170 cm^-1^ (Fig. S3). When the Raman signal intensity shows a linear dependence on the material density, a calibration coefficient (k) can be calculated. The arbitrary Raman intensity CCD readout counts were converted to absolute surface material density of poly-P by means of the calibration coefficient (k) as detailed earlier ^36^. A sodium polyphosphate standard solution (0.5 μL) with a concentration of 0.1 mg P mL^-1^ was mounted and air dried on the CaF_2_ Raman window. The dried poly-P droplet occupied an area of about 0.3 mm^2^ (Fig. S4), corresponding to a material density about 0.16 μg mm^-2^. The averaged material density can be written as;

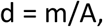

where d is material density (g/μm^2^), m is amount of material (g) and A is the scanned area (μm^2^). Furthermore,

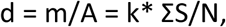

where k is the calibration coefficient (g/μm^2^/count), ΣS is the cumulative analyte Raman signal strength (counts) and N is the total spectra sampling points in the designated map area. Rearrangement gives k:

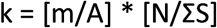

The area of the mapped droplet was calculated by ImageJ software and was estimated to be 324,473 μm^2^ (Fig. S4) ^37^. The total mapped area contained 1452 individual spectra. The cumulative poly-P Raman signal strength at the Raman marker wavenumber (1170 cm^-1^) recorded for the scanned area was 450,042 CCD counts. This gives a k for poly-P for this particular set of conditions of 4.98 ± 0.2 * 10^-16^ g P μm^-2^ count^-1^ (n = 6).

Assuming that *T. elongata* cells are rods with hemispherical ends, the two-dimensional area occupied by cells on the Raman slide was estimated using ImageJ software. The average estimated area occupied by *T. elongata* cells on CaF_2_ Raman slides was 3.79 μm^2^ (n = 100 single cells). *T. elongata* cell counts were conducted as detailed earlier ^38^. *T. elongata* cell number in the culture (DAPI stained cell counts) was determined to be 7.9 ± 1.7 *10^7^ cells ml^-1^. These values were used for the Raman-based estimation of the fraction of phosphorus being assimilated in the form of poly-P within *T. elongata* cells under P-uptake conditions. Cell volumes were calculated from fluorescence images of *Tetrasphaera* and *Ca*. Accumulibacter (n = 100 cells) as described earlier in ^39^. P accumulation per *T. elongata* cell was validated using bulk liquid P quantification at key stages of the P-uptake release experiment (Supplementary text – 2).

Standard of sodium polyphosphate (CAS # - 68915–31–1) was obtained from Sigma Aldrich, UK. Adenosine 5'-triphosphate (ATP) disodium salt was purchased from Fluka (Germany) and adenosine 5'-monophosphate (AMP) sodium salt from Boehringer Mannheim GMBH (Germany).

### Analysis of PHA and glycogen

Absolute quantification of PHA and glycogen was conducted using the same approach as for poly-P. Poly-3-hydroxybutyric acid-co-3-hydroxyvaleric acid dissolved in CHCl_3_ and glycogen dissolved in water were mounted on CaF_2_ Raman windows in a range of densities to estimate their calibration coefficients. PHB-co-HV and glycogen droplets were Raman mapped as described above and k-values were determined to be 1.04 ± 0.1 * 10^-14^ g C μm^-2^counts^-1^ and 2.37 ± 0.2 * 10^-15^ g C μm^-2^ counts^-1^, respectively. PHAs produce characteristic Raman peaks at 432 cm^-1^, 840 cm^-1^, and 1726 cm^-1^, respectively, that are attributed to δ (C-C) skeletal deformations and v(C=O) stretching vibrations ^40^, whereas glycogen produces strong characteristic Raman peaks within the regions of 478 – 484 cm^-1^ and 840 – 860 cm^-1^, which are attributed to C-C skeletal deformation and CC skeletal stretch, respectively ^39^ (Fig. S1). For PHA and glycogen, Raman markers at 1726 cm^-1^ and 481 cm^-1^ respectively, were used for all quantifications. Raman calibrations for various material densities of PHA and glycogen are shown in Fig. S5.

Standard glycogen sourced from oyster (CAS # - 9005–79–2) and poly (3-hydroxybutyric acid-co-3-hydroxyvaleric acid) (CAS # - 80181–31–3) were obtained from Sigma Aldrich, UK.

### Combined FISH-Raman analysis of activated sludge

FISH analyses were conducted as described earlier ^32^, but without the addition of sodium dodecyl sulfate (SDS) in the final washing buffer in order to minimize loss of intracellular storage compounds and to minimize biomass loss from CaF_2_ Raman slides during FISH washing procedure. The FISH probes applied in this study were the *Ca*. Accumulibacter-specific PAO651 probe ^8^, the *Tetrasphaera-specific* probe Actino658 ^11^ and the general EUBmix probe set (EUB 338, EUB 338 II, and EUB 338 III) ^41^ targeting most bacteria. *Ca*. Accumulibacter is routinely detected with the PAOmix probe set (including PAO462, PAO651, and PAO846) ^8^, however, it was recently shown that the PAO651 probe alone gives better specificity for the genus without a substantial sacrifice in coverage ^42^. In the 8 EBPR plants investigated here, the biovolume quantified by the PAOmix probe set was 5–15% greater than with the PAO651 probe with the difference almost exclusively made up by *Propionivibrio* targeted by the probe Prop207 ^42^ (Supplementary text-3). *Propionivibrio* is supposed not to be a PAO ^42^ and our Raman investigations confirmed that it did not contain polyp *in situ*. Several probes are available to target sub-groups of the *Tetrasphaera* genus in activated sludge ^11, 37^ – however our recent study revealed low diversity of the genus in Danish systems^12^ and the Actino658 probe provides the most specific coverage of the dominant member of the genus. In addition, *in silico* analyses of the specificity of the sub-group probes shows that they cannot any longer be regarded as specific (S. McIlroy, unpublished). The 5' ends of the oligonucleotide probes were labelled with 5(6)-carboxyfluorescein-N-hydroxysuccinimide ester (FLUOS) or with the sulfoindocyanine dyes (Cy3) (Thermo-Fisher Scientific, Germany). The FISH signal was observed without mounting media using the in-built fluorescence microscope of the Raman system. Labelled single cells of interest were identified and marked for Raman analysis using the LabSpec 6.4 software. As the fluorescent signal from the Cy3 label interferes with the Raman signal the former was subsequently bleached as previously described ^43^. To achieve this, the 532.17 nm Raman laser, set at the intensity used for sample analysis, was shone on the analysed area of the sample for 5 min (Fig. S6). The Raman signal from FISH-identified-cells of interest was then acquired with the same settings as described for the pure culture analyses.

### Quantitative FISH analysis of full-scale sludge samples

Quantitative FISH (qFISH) values of PAO populations were calculated as a percentage area of the total biovolume stained with the EUBmix probe set ^44, 40^ that also hybridized with the specific probe. qFISH analyses were based on 30 fields of view taken at 630 x magnification using the Daime image analysis software ^45^. Microscopy was performed with a White Light Laser Confocal Microscope (Leica TCS SP8 X, Leica Microsystems, Kista, Sweden). Quantitative FISH was conducted on fixed samples from the aeration tanks of all EBPR plants investigated. The qFISH results were similar for samples obtained from other process tanks due to high recirculation rates (data not shown).

### Analysis of P and PHA leakage from cells due to cell fixation and the FISH procedure

Potential loss of intracellular polymers during fixation of cells (in either PFA or ethanol) and the FISH procedure (including 4 h incubations in formamide hybridization buffer at 46°C) were investigated, as these steps were performed prior to Raman analyses. To test this, the Gram-positive PAO *T. elongata* and the Gram-negative organism *A. junii*, harvested at growth stages at which the intracellular poly-P content was high, were used. *A. junii* was chosen because no *Ca*. Accumulibacter monoculture has been isolated to date and *A. junii* cells are known to take up large quantities of poly-P ^46^. *A. junii* was grown in a modified Fuhs and Chen semi-defined medium ^47^ containing extra ortho-P. The composition of the medium was as follows (per Litre): Sodium acetate, 5 g; (NH_4_)_2_SO_4_ ,2 g; MgSO_4_.7H_2_O, 0.5 g; KH_2_PO_4_, 0.25 g; CaCl_2_.2H_2_O, 0.2 g; casamino acids, 0.6 g. The culture was grown at 30°C for 96 h. The Raman signal for poly-P accumulation in unfixed *A. junii* cells was highest on day 4. Aliquots of the two monocultures, *T. elongata* and *A. junii*, with a high poly-P content were homogenized and fixed. The fixed cells were subjected to the standard FISH procedure using the general EUB338 mix probes. Both fixed and unfixed preparations of the *T. elongata* and *A. junii* cells, with and without FISH, were then analyzed by Raman microspectroscopy for their intracellular poly-P contents (n=100 cells in each instance). For assessing PHA loss during cell fixation and storage, pure cultures of *Bacillus subtilis* and *Pseudomonas aeruginosa* were used as representatives of Gram-positive and Gram-negative, PHA producing, model organisms. *B. subtilis* and *P.aeruginosa* were grown aerobically in media known to give a high PHA yield ^48, 49^.

### Chemical analyses

Analysis of bulk ortho-P levels was carried out using the ammonium molybdate based colorimetric standard method ISO 6878:2004. The cultures were centrifuged at 4,000 x g and the supernatant was filtered through 0.22 μm polyvinylidene fluoride (PVDF) filters (Millipore, UK) and the filtrate was used for bulk ortho-P determinations. The chemical oxygen demand (COD) of the samples was determined using the acidified dichromate-based colorimetric commercial kit LCK-414 (HACH-Lange, UK). The total P content of the activated sludge samples was analyzed by inductively coupled plasma – atomic emission spectroscopy (ICP-AES), as described earlier ^50^.

## Results and Discussion

### Poly-P accumulation in *T. elongata* during feed-famine cycling

*T. elongata* is one of the few known PAO that exist in pure culture and it was thus selected to test for qualitative and quantitative assessment of its intracellular storage polymers using Raman microspectroscopy. However, it should be noted that *T. elongata* is closely related but not identical to *Tetrasphaera* strains found in full-scale WWTPs ^37^. *T. elongata* cultures showed the typical ortho-P uptake/release patterns of a PAO during sequential feed-famine cycling (Fig. 1A). The increase of bulk liquid ortho-P concentration during the anaerobic “feed” conditions was coupled to uptake of organic substrate from the bulk medium, reflected by the decrease of COD. The drop in ortho-P levels in the bulk medium during the aerobic conditions was reflected by a strong increase of poly-P in *T. elongata* cells (see below).

**Figure 1:**
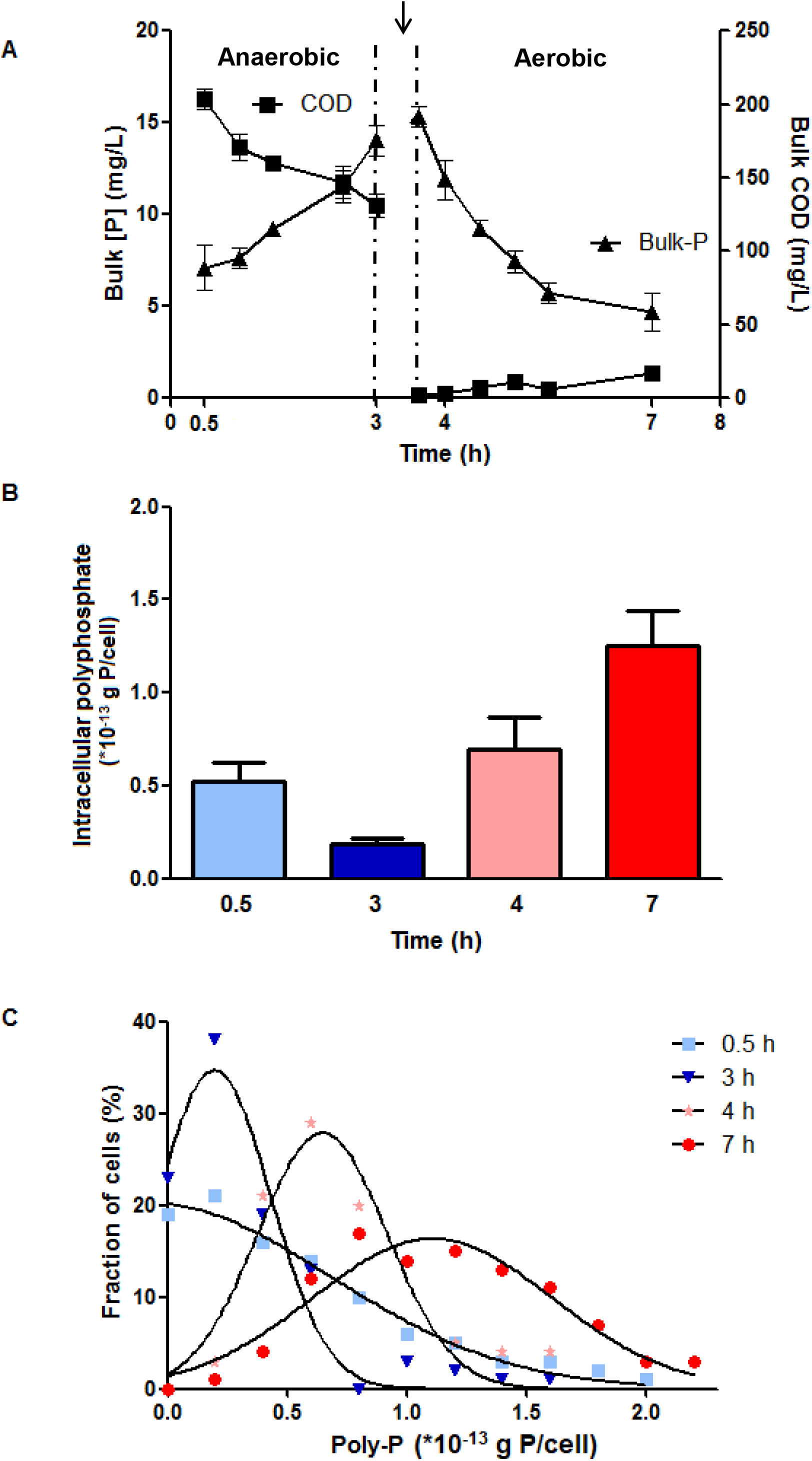
(A) Bulk medium ortho-P concentration and COD changes during P-uptake/P-release experiments using a *T. elongata* pure culture. The patterns reflect substrate uptake during the anaerobic feed phasecoupled to P release, and P uptake during the subsequent aerobic famine phase. The arrow indicates the point at which cells were washed and the medium made aerobic. (B) Average changes of the cellular poly-P content as determined by Raman microspectroscopy on unfixed cells during the P-uptake/release experiment. (C) Distribution pattern of *T. elongata* cells based on their intracellular poly-P content during various time points of the P-uptake/release experiment (n = 100 individual *T. elongata* cells in each instance, mean ± SD in error bars).

Raman microspectoscopy was applied to measure the intracellular poly-P storage at different stages of the feed-famine cycling. As expected, the average poly-P content decreased during anaerobic conditions and increased during aerobic famine conditions. (Fig. 1B). The changes in the bulk P concentration and the dynamics of the intracellular P throughout the experiment is corroborating evidence that the P changes in the bulk medium were due to the release and uptake of P by *T. elongata*. The P-content of individual cells showed a broad distribution (Fig. 1C) with maximum values around 2*10^-13^ g P cell^-1^ at the end of the aerobic stage. Under anaerobic conditions, a large fraction of cells had no measurable poly-P. The average cellular P content at the end of the anaerobic (3 h) and aerobic (7 h) stages was 0.19*10^-13^ g P cell^-1^ and 1.2*10^-13^ g P cell^-1^, respectively (calculations in Supplementary text - 4). The broad distribution of intracellular poly-P contents in the aerobic stage shows population heterogeneity in the capability to store poly-P, perhaps due to variation in growth stages or different strains covered by the broad probe. Based on these calculations, the average intracellular poly-P during the aerobic phase corresponded to 8.0 mg P l^-1^. The difference in this value to the same estimation from bulk ortho-P levels of 9.3 mg P l^-1^ is likely explained by the incorporation of some of the ortho-P into the biomass during growth.

These results demonstrate the utility of Raman microspectroscopy for the direct quantification of intracellular poly-P and the analysis of its storage dynamics under *in situ* conditions. Previous studies have studied such dynamics only in a relative manner, with the P-content normalized to the biomass using a biomass marker (amide I at 1660 cm^-1^) ^30^. Other qualitative imaging-based assessments of poly-P inclusions such as DAPI staining ^51^ have their inherent disadvantages due to their inability to quantify the levels of poly-P accumulation and their non-specific binding to other cellular components such as lipids and nucleic acids ^20, 52^. Surprisingly and inconsistent with the metabolic model inferred from comparative genomics, we could not detect glycogen in *T. elongata* by Raman microspectroscopy at any stage of the P-uptake/release experiments (Fig. S7).

### Dynamics of storage compounds in *Ca*. Accumulibacter and *Tetrasphaera* in activated sludge during feed-famine cycling experiment

Initially, potential loss of intracellular polymers during cell fixation and FISH was assessed with cultures of representative Gram-negative and Gram-positive bacteria. Cell fixation and FISH lead to a loss of up to 20% of the overall poly-P Raman signal for both types of bacteria (Fig. S8. Similarly, for PHAs, the Raman signal loss following fixation and FISH amounted to a maximum of 12%, for both Gram-negative and Gram-positive bacteria. Therefore, the applied FISH-Raman method for quantitative analyses of FISH-identified cells will slightly underestimate the actual amount of these storage compounds. An average loss of 12% and 8% for poly-P and PHA, respectively, have been corrected for in the analyses and mass balances presented in this study.

In the next step, several batch P-cycling experiments with fresh activated sludge from Aalborg West WWTP were carried out to provide insights into the *in situ* dynamics of intracellular polymers in FISH probe-defined *Ca*. Accumulibacter and *Tetrasphaera* (Fig 2). During the anaerobic feed phase, most of the *Ca*. Accumulibacter and *Tetrasphaera* cells contained little intracellular poly-P (on average 0.48*10^-13^ g P cell^-1^ and 0.27*10^-13^ g P cell^-1^, respectively – Fig. 3B and 1B). During the aerobic famine phase, most cells of both species contained several fold higher levels of poly-P than in the anaerobic phase (average of 6.46*10^-13^ g P cell^-1^ and 1.77*10^-13^ g P cell^-1^, respectively). The fluctuations of intracellular poly-P contents corroborated well with the dynamics of the bulk ortho-P concentrations observed during the incubation (Fig. 3A, 3B and 3C). These lab-scale P uptake/release experiments demonstrated that in this activated sludge *Ca*. Accumulibacter cells were capable of storing at least three times as much poly-P per cell compared to *Tetrasphaera*. However, as the cell volumes of the two PAO types are very different, with *Ca*. Accumulibacter nearly three times larger than *Tetrasphaera* (9.2±2.4 μm^3^ and 2.9±1.1 μm^3^), both PAO clades had very similar volumetric average maximum P-contents of 0.70 * 10^-13^ g P μm^-3^ and 0.68 * 10^-13^ g P μm^-3^, respectively. Consequently, the maximum P-storage capacity of both PAO clades per biovolume was highly similar in this WWTP. Recently, it was shown that some P-uptake (without demonstrated poly-P formation) may occur at anaerobic conditions of P-cycling experiments of *Tetrasphaera* enriched lab-scale cultures ^53^. This effect, however, was not observed throughout this study, either in anaerobic stages of pure culture experiments employing *T. elongata* monocultures, or *Tetrasphaera* cells found *in situ* in full-scale EBPR plants (see later).

**Figure 2:**
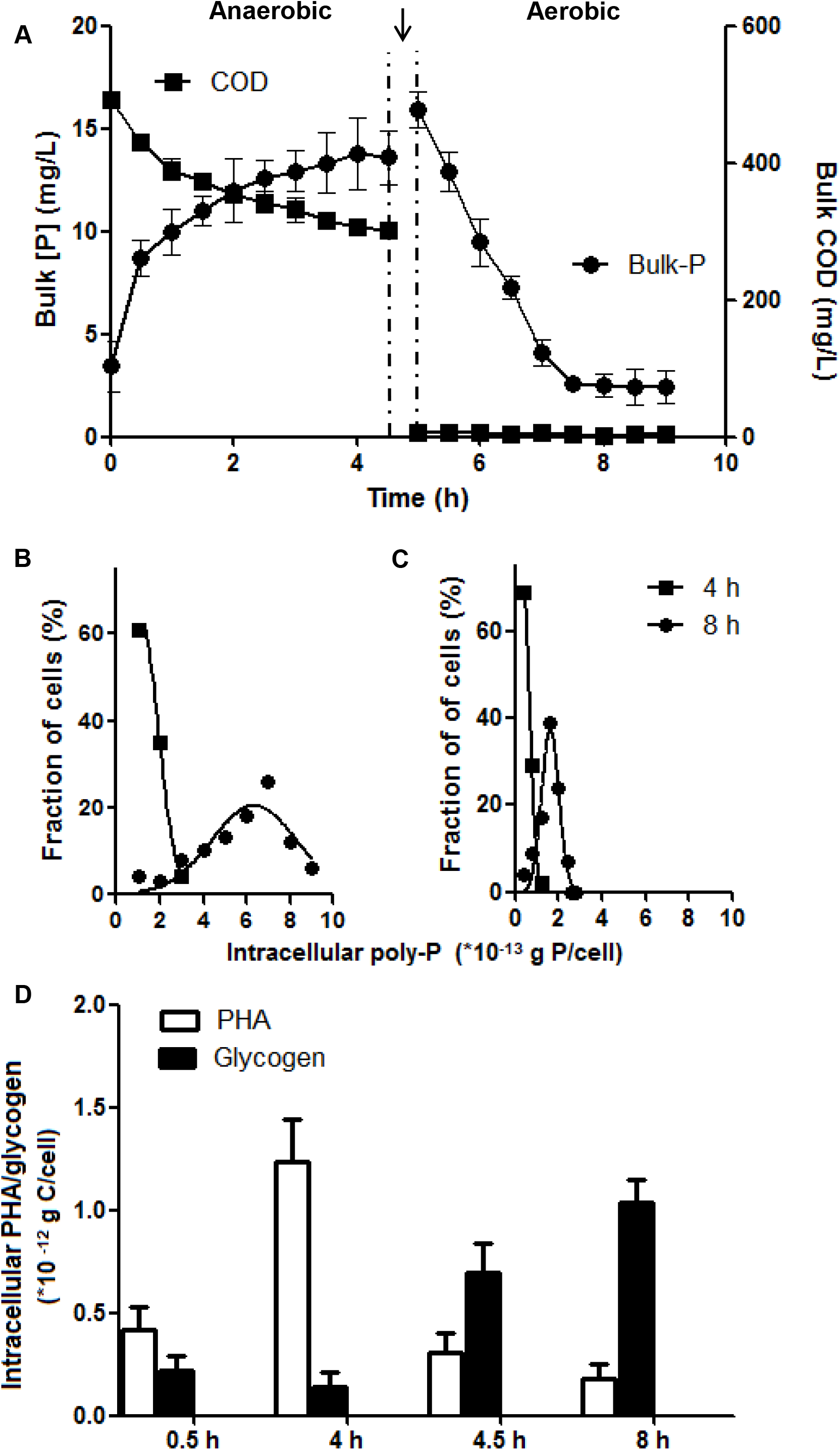
Dynamic of intracellular polymers in probe-defined *Ca*. Accumulibacter and *Tetrasphaera* in lab–scale experiments with activated sludge from Aalborg West full–scale EBPR plant. A) Bulk medium concentrations of ortho-P and COD changes during 4 h anaerobic and 8 h aerobic time course experiments. The arrow indicates the point at which the mixed biomass was washed and the medium was made aerobic. B) Intracellular poly-P contents of *Ca*. Accumulibacter cells. C) Intracellular poly-P contents of *Tetrasphaera* cells (n = 100 individual probe defined cells in each instance, mean ± SD in error bars). (D)Intracellular changes in glycogen and PHA content in *Ca*. Accumulibacter cells.

**Figure 3:**
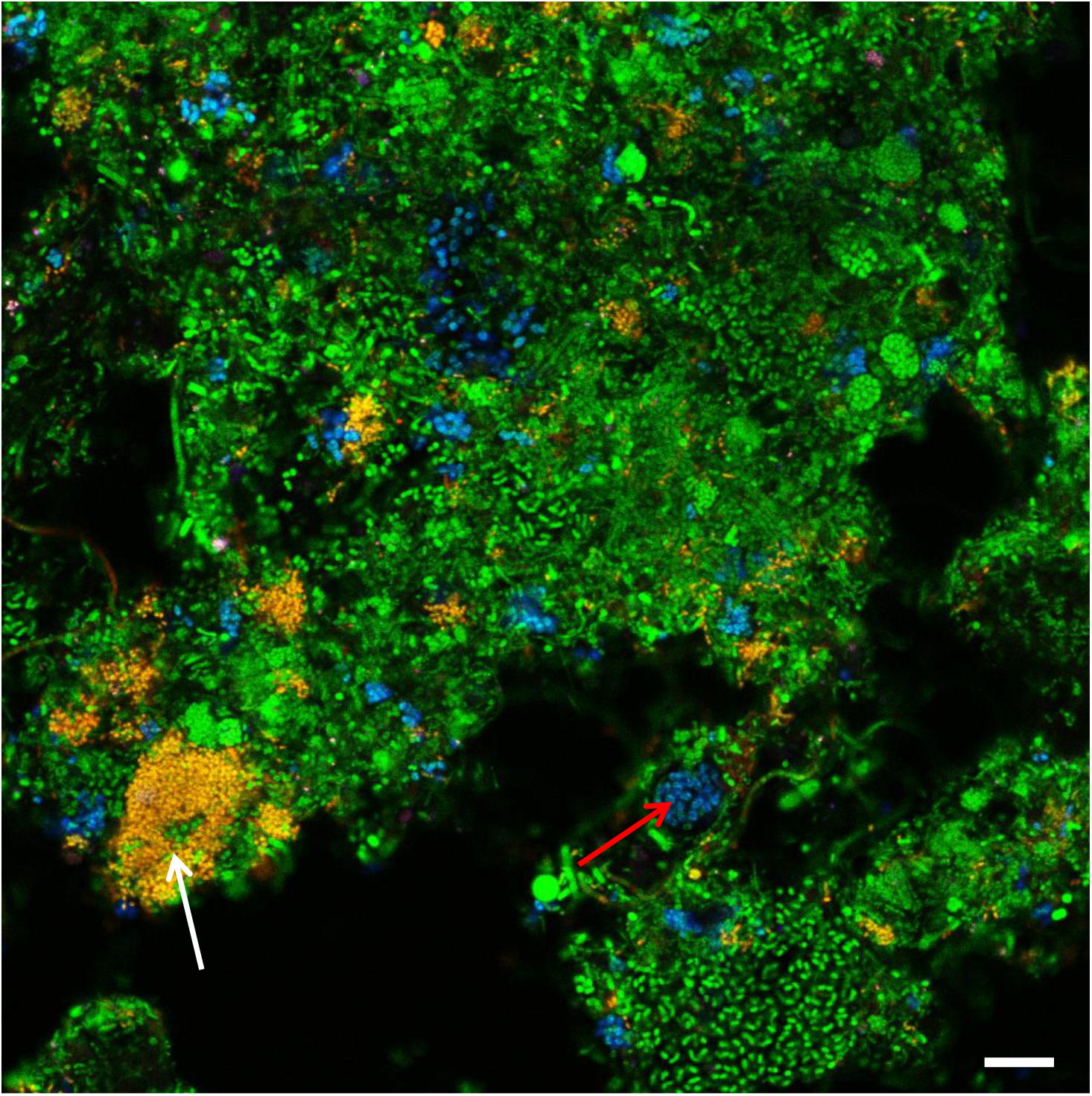
Composite FISH image of the PAO in the Aalborg West WWTP. *Tetrasphaera* appears yellow/orange (overlap of hybridization signals from probe Actino658 (Cy3, red), and the EUB probe mix (FLUOS, green). *Ca*. Accumulibacter appears cyan (overlap of hybridization signals from probe PAO651 (Cy5, blue) and the EUB probe mix (FLUOS, green) (Scale bar, 10 μm). White and red arrows indicate *Tetrasphaera* and *Ca*. Accumulibacter cells, respectively.

In accordance with the generally accepted metabolic model for PAO ^17, 7^, intracellular PHA and glycogen in *Ca*. Accumulibacter showed a marked difference in Raman signal intensities between the anaerobic and aerobic phase (Fig. 3D). PHA accumulated during the anaerobic feed phase and was consumed during the aerobic famine phase with glycogen showing the opposite pattern. Consistent with the pure culture experiments with *T. elongata* and previous literature data ^11, 18, 19^, probe-defined *Tetrasphaera* cells in the activated sludge did not show Raman signatures for PHAs during the P-cycling experiments. Surprisingly but consistent with the pure culture data (Fig. S7), no glycogen was detected in probe-defined *Tetrasphaera* in activated sludge. Furthermore, no other potential storage compound detectable by Raman spectroscopy was observed in *T. elongata* (Fig. S7) or in probe-defined *Tetrasphaera* in activated sludge. Possibly, *Tetrasphaera* relies under anaerobic conditions on hydrolysis of poly-P in addition to possible fermentation of glucose or amino acids for anaerobic growth, and/or storage of these for subsequent growth and replenishment of poly-P during aerobic conditions. Another possibility is that the presence of glycogen in *Tetrasphaera*, as reported in other studies, is wrong. Hitherto, no glycogen specific analysis method has been employed to quantify the polymer in its native state within single bacterial cells, including *Tetrasphaera*. The glucose monomeric units measured after acid hydrolysis of cells were inferred to originate from glycogen stored in cells in all of the previous studies ^18^. This obviously is not optimal due to the ubiquity of glucose and other similar sugars in the intracellular milieu, even when the cells are incapable of storing glycogen. The strength of the current FISH-Raman method is that it does not rely on such inferences and that it can directly measure the polymer in question in its native state at a single-cell level. It has been shown that *T. elongata* can grow by fermentation ^19^, and probe-defined *Tetrasphaera* in full-scale EBPR plants likely ferment glucose after several of days anaerobic conditions ^54^, so fermentation is likely a key metabolic feature in tandem with poly-P formation/degradation in their successful competition to many other microbes in EBPR plants.

### *In situ* quantification of storage polymers in *Ca*. Accumulibacter and *Tetrasphaera* in full-scale EBPR plants

*Ca*. Accumulibacter has generally been considered as the most important PAO in EBPR plants world-wide ^7^. However, recent investigations have indicated that *Tetrasphaera* are more abundant in many full-scale plants ^12^, but abundance data alone are insufficient to judge the importance of a PAO for the EBPR process. As the ecophysiologies of these two PAO clades appear to be very different, it is of primary importance to know what their actual relative contributions to P removal in full-scale EBPR systems are, in order to lay the foundation for knowledge-based optimization of plant design and operation.

In the current study, this key question was tackled by analyzing eight full-scale EBPR plants representing two different designs (recirculating and alternating) that all have had good and stable operation for several years ^12^ (Table S1 and S2). Bulk concentration of ortho-P in the aerobic tank of all plants was 0.2–0.6 mg P l^-1^, which was well below the maximum tolerated effluent concentration of 1.0 mg P l^-1^ (see Table S1). In the different tanks of these EBPR plants, the biovolume of *Tetrasphaera* and *Ca*. Accumulibacter and the amount of intracellular poly-P, PHA, and glycogen in these PAO was measured by quantitative FISH and FISH-Raman microspectroscopy, respectively. qFISH analyses showed that *Ca*. Accumulibacter and *Tetrasphaera* were present in all plants in abundances of 1.2 to 6.6% (relative biovolume of the target population to the total biovolume of all FISH-detectable cells). *Tetrasphaera* outnumbered *Ca*. Accumulibacter in 5 of the 8 plants investigated (Fig. 3, Table 1). In accordance with the result from the batch experiments (Fig. 3), and the current EBPR model, the level of intracellular poly-P in both PAO was lowest in the anaerobic tanks and highest in the aerobic tanks. As predicted by the EBPR model, the PHA levels were also dynamic in *Ca*. Accumulibacter with highest levels in the anaerobic tank and lowest under oxic conditions, reflecting uptake of short-chain fatty acids in the anaerobic tank and growth and respiration in the oxic tank. Likewise, the intracellular glycogen content in *Accumulibacter* was lowest in the anaerobic tanks and highest in the aerobic tanks, reflecting the production of reducing power for PHA formation under anaerobic conditions and replenishment of glycogen from PHA in the presence of oxygen, respectively. Consistent with the batch experiments with sludge from a single plant, PHA and glycogen were not observed in *Tetrasphaera in situ* in any of the full-scale EBPR plants (to enhance clarity, the results for two typical plants are shown in Fig. 4 while the remaining six are shown in Fig. S9). The stoichiometry of the formation/degradation of the intracellular storage compounds for *Ca*. Accumulibacter cells that we observed using the Raman-FISH based single-cell method for the full-scale plants are fully consistent with bulk data reported previously from lab-scale studies using enriched biomass of this PAO. For example, the molar ratio of glycogen formation/PHA degradation (C/C) of 0.3 – 0.4 that we measured in the aerobic tanks of the EBPR plants, fully matches data obtained in lab-experiments with activated sludge enriched in *Ca*. Accumulibacter ^55, 9^. These consistencies provide additional strong support for the suitability of FISH-Raman microspectroscopy to quantitatively follow storage compound dynamics *in situ* in FISH probe-defined taxa.

**Figure 4:**
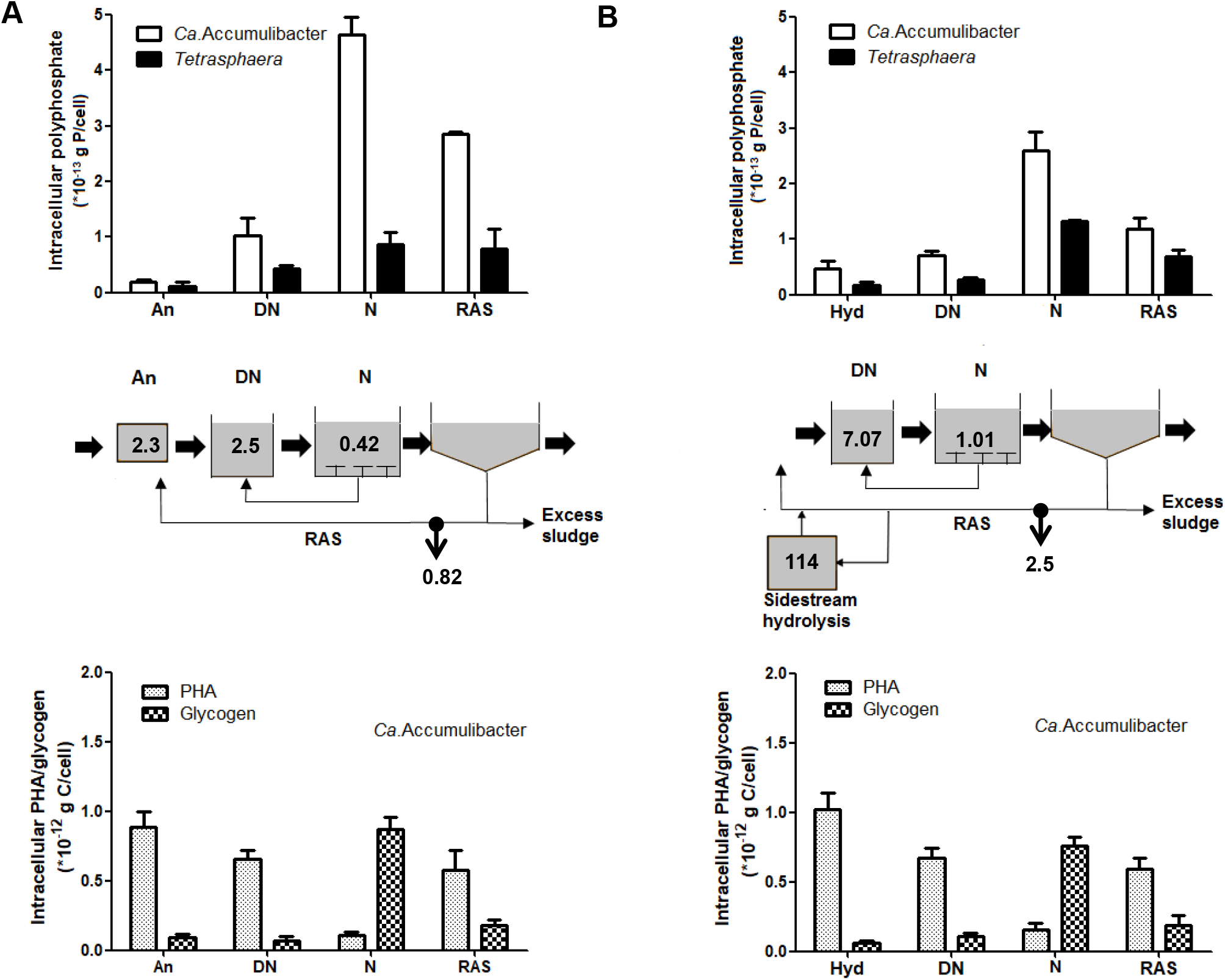
Dynamics of intracellular poly-P and PHA/glycogen in *Ca*. Accumulibacter and *Tetrasphaera* in the different process tanks in HjØrring (A) and Aalborg West (B) EBPR plants. Intracellular poly-P and intracellular PHA/glycogen are shown for *Ca*. Accumulibacter while no PHA or glycogen was found in *Tetrasphaera* and thus not shown. An, Hyd, DN, N and RAS denotes anaerobic, sidestream hydrolysis, denitrification, nitrification (aeration) tanks and return activated sludge, respectively. The numbers in each stage of the figures indicate the bulk ortho-P concentrations in mg P L^-1^, mean ± SD in error bars, n = 100 individual probe defined random cells in each instance).

**Table – 1:** A summary of all WWTPs investigated, the individual P removal contributions from the two main PAOs, *Ca*. Accumulibacter and *Tetrasphaera* spp. (*the individual percentage contribution from each PAO to the overall P removal process in each WWTP is denoted within parentheses, Alt – alternating operation, Rec – recirculation, An – anaerobic tank, Hyd – sidestream hydrolysis).

| WWTP location | Design type and presence of anaerobic tank and/or sidestream hydrolysis | qFISH Biovolume fraction (%) |  | Poly-P in the N tank |  | Ortho -P in the N tank (mgP L<sup>-1</sup>) | Normalized Total P (mgP gSS<sup>-1</sup>) | Fraction *Ca.* Accumulibacter (mgP gSS<sup>-1</sup>)<br>*contribution to overall P removal (%) | Fraction *Tetrasphaera* (mgP gSS<sup>-1</sup>)<br>* contribution to overall P removal (%) |
| --- | --- | --- | --- | --- | --- | --- | --- | --- | --- |
|  |  | *Ca.* Accumulibacter | *Tetrasphaera* | *Ca.* Accumulibacter (*10<sup>-13</sup> gP cell<sup>-1</sup>) | Tetrasphaera (*10<sup>-13</sup> gP cell<sup>-1</sup>) |  |  |  |  |
| Hjørring | Rec (An) | 1.20 ± 0.5 | 6.50 ± 0.6 | 4.63 ± 0.32 | 0.86 ± 0.23 | 0.43 ± 0.041 | 21.72 ± 2.7 | 2.08 ± 0.1 (9.5) | 6.72 ± 0.51 (30.9) |
| Randers | Rec (Hyd) | 2.34 ± 0.7 | 6.64 ± 0.2 | 4.08 ± 0.41 | 1.35 ± 0.15 | 0.45 ± 0.26 | 22.74 ± 0.48 | 3.4 ± 0.39 (14.9) | 10.57 ± 0.34 (46.5) |
| Viby | Rec (Hyd) | 2.72 ± 1.3 | 2.55 ± 0.7 | 3.14 ± 0.39 | 0.98 ± 0.29 | 0.57 ± 0.22 | 24.81 ± 1.2 | 3.13 ± 0.22 (12.6) | 2.91 ± 0.21 (11.7) |
| Bjerringbro | Alt (An) | 4.96 ± 1.4 | 3.53 ± 0.2 | 4.02 ± 0.39 | 1.25 ± 0.34 | 0.55 ± 0.01 | 17.75 ± 0.92 | 7.35 ± 0.44 (41.4) | 5.19 ± 0.29 (29.2) |
| Aalborg East | Alt (Hyd) | 1.60 ± 0.6 | 2.20 ± 0.9 | 4.63 ± 0.49 | 1.33 ± 0.17 | 0.53 ± 0.07 | 22.51 ± 3.1 | 2.77 ± 0.72 (12.3) | 3.8 ± 0.37 (16.9) |
| Aalborg West | Alt (Hyd) | 3.60 ± 1.6 | 3.10 ± 1.6 | 3.68 ± 0.39 | 1.39 ± 0.2 | 1.01 ± 0.21 | 37.52 ± 2.9 | 5.04 ± 0.28 (13.4) | 5.17 ± 0.33 (13.8) |
| Egaa | Alt (An/Hyd) | 1.71 ± 0.4 | 2.04 ± 0.8 | 4.22 ± 0.36 | 1.47 ± 0.41 | 0.28 ± 0.02 | 17.33 ± 0.35 | 2.66 ± 0.24 (15.4) | 3.47 ± 0.39 (20) |
| Åby | Alt (An/Hyd) | 2.53 ± 1.4 | 3.84 ± 1.1 | 3.83 ± 0.18 | 1.29 ± 0.62 | 0.33 ± 0.12 | 14.84 ± 0.74 | 3.55 ± 0.64 (23.9) | 5.70 ± 0.47 (38.4) |

Interestingly, in none of the eight WWTP the *Ca*. Accumulibacter stored *in situ* under aerobic conditions as much poly-P per cell than in the lab-scale incubation experiment with activated sludge from the Aalborg West WWTP (that was also included in the eight WWTP) and plenty of organic substrate in the anaerobic phase. While 6.5 * 10^-13^ g P cell^-1^ were measured in the lab-scale experiments, the highest recorded *in situ* value for *Ca*. Accumulibacter was 4.6 * 10^-13^g P cell^-1^ in the Hjørring WWTP and the lowest *in situ* value was 3.1 * 10^-13^g P cell^-1^ in the Viby plant. Similarly, the poly-P content of the *Tetrasphaera* cells in all plants was lower than in the lab-scale experiment with activated sludge (1.77* 10^-13^g) with the highest value (1.5 * 10^-13^g P cell^-1^) recorded in the Egaa plant and the lowest value of 0.89 * 10^-13^ g P cell^-1^ measured in the Viby plant (Fig. 4 and Fig. S9. The level of PHA uptake for *Ca*. Accumulibacter cells was similar in all plants in the anaerobic tanks (0.9–1.4 * 10^-12^g C cell^-1^), corresponding well to the maximum capacity seen in the lab-scale activated sludge experiment with surplus substrate of 1.3 * 10^-12^g C cell^-1^ (Fig. 3). PHA was subsequently consumed by *Ca*. Accumulibacter in the anoxic denitrification and aerobic tanks. In some plants this process occurred primarily under denitrifying conditions (Fig. S9) indicating differences in the presence of denitrifying members of this genus ^56, 18^. Consistently, the glycogen level of *Ca*. Accumulibacter cells was highest in the aerobic tanks (0.7–0.9 * 10^-12^ g C cell^-1^) and always slightly lower than in the lab-scale experiment (Fig. 3) (1.1 * 10^-12^ g C cell^-1^). The absence of glycogen in detectable quantities throughout P-uptake/release experiments in *T. elongata* monoculture experiments, and *in situ* in *Tetrasphaera* cells in activated sludge samples (in contrast to *Ca*. Accumulibacter) demonstrates that glycogen did not act as a carbon reserve during P-cycling in *Tetrasphaera* (limit of detection for analytes in this study was determined as described in Supplementary text – 5). The currently accepted standard model says that glycogen should act as the carbon storage polymer, providing energy and reducing equivalents to the uptake of P and storage as poly-P. This work does not conform to this notion, but rather suggests that other cellular carbon reserves (such as fermentation products or amino acids) may provide the reducing equivalents needed.

### *Tetrasphaera* is more important than *Ca*. Accumulibacter for P-removal

Based on the data obtained from FISH-Raman-microspectroscopy and from qFISH, the relative contribution of *Ca*. Accumulibacter and members of the genus *Tetrasphaera* to the P-removal in the eight full-scale EBPR WWTP was calculated. The following P pools were considered: poly-P stored in *Ca*. Accumulibacter and *Tetrasphaera*, P incorporated into assimilated biomass (nucleic acids, cell membranes etc.), poly-P in unknown PAO, and P bound in chemical precipitates (primarily iron, calcium, and aluminium) with the latter two pools being treated as one as no approach for their differentiation is available (calculations details are provided in the supplementary material). *Ca*. Accumulibacter and *Tetrasphaera* were both important in all plants and contained together 24 – 70% of the total P. In 6 of the 8 plants the contribution of *Tetrasphaera* to the total P-removal was higher than that of *Ca*. Accumulibacter (Fig. 5; Table 1). Our data also demonstrate that there is a relatively high fraction of removed P in all plants that could either represent chemically precipitated P or be assigned to the activity of yet undescribed PAO (see P mass balance calculations in Supplementary text-6). Future studies might want to use a combination of the Raman microspectroscopy approach developed here, the recently emerging Raman cell sorting techniques ^57^, and single cell genomics to hunt in a targeted manner for such not yet discovered PAO.

**Figure 5:**
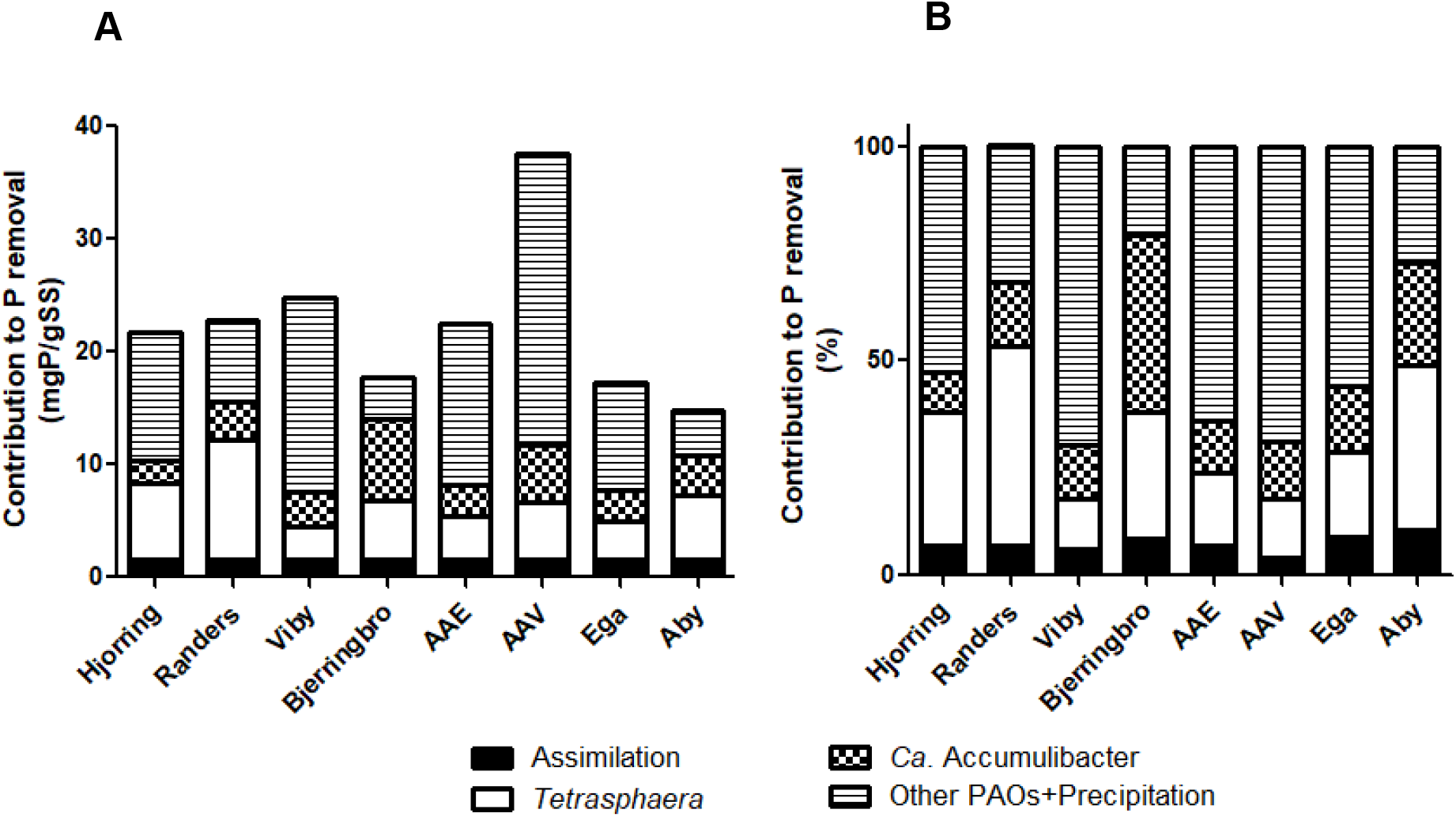
(A) Absolute distribution of P pools in activated sludge in the aeration tanks of eight full-scale EBPR plants, and (B) the percentage of total P in the different pools.

In the next few years many EBPR plants will be build world-wide in order to improve effluent quality of WWTP in a cost-effective and sustainable way and for the recovery of P from wastewater. This study showed that the long-held view that members of *Ca*. Accumulibacter, and their “classical PAO” physiology are driving EBPR must be significantly revised. Our study demonstrates for the first time that the “non-classical” fermentative PAO *Tetrasphaera* are contributing in the majority of the analysed plants more to EBPR than *Ca*. Accumulibacter and future studies are now urgently needed to test this on a global scale. Furthermore, it will be important to determine the affinity (half saturation coefficient for ortho-P) of both PAO clades as promotion of the growth of high affinity PAO will be the desirable in plants with low P-effluent standards. The unexpected insights obtained in this study became possible via combining quantitative FISH and quantitative Raman microspectroscopy for absolute quantification of intracellular storage compounds in FISH-identified PAO taxa. This single-cell *in situ* approach enables to quantitatively determine the contribution of individual microbial taxa to storage compound formation in complex microbial communities and thus offers new opportunities for an in depth functional understanding of EBPR and all other microbial ecosystems in which such processes are important.

## Acknowledgements

The study was supported by Innovation Fund Denmark (ReCoverP, 4106–00014B and NomiGas, 1305–00018B). K. Hansen, J. Dencker, A. Boisen and K. Reitzel are acknowledged for analyses of total P. M.S. and M.W. were also supported by the European Research Council Advanced Grant project NITRICARE 294343 (to M.W.).

